# Pasting behaviour of Australian native grain flours: effects of particle size, flour blending, and temperature

**DOI:** 10.64898/2026.09.29.755466

**Authors:** Farkhondeh Abedi, Claudia Keitel, Ali Khoddami, Thomas H. Roberts

**Affiliations:** School of Life and Environmental Sciences, University of Sydney, Camperdown NSW 2050, Australia; Sydney Institute of Agriculture, University of Sydney, Eveleigh, NSW 2015, Australia

**Keywords:** Australian native grains, flour functionality, pasting properties, Rapid Visco Analyser, particle size, composite flour

## Abstract

Knowledge of the influence of particle size, blending, and processing temperature on flour functionality is critical for the utilisation of emerging grains. This study investigated the flour pasting behaviour of four Australian native grains—Button Grass, Curly Mitchell Grass, Native Millet, and Weeping Grass—in relation to particle size, wheat flour substitution at various levels (0–30%), and elevated RVA temperatures (95–140 °C). The pasting properties of flour fractions (fine, medium, and wholegrain) were analysed using a Rapid Visco Analyser, and water absorption capacity was also determined. Particle size effects were species-specific. Native Millet and wheat had higher viscosity in the finer fractions, whereas Weeping Grass exhibited maximum viscosity in the medium fraction. Button Grass and Curly Mitchell Grass showed limited sensitivity to particle size, indicating a stronger influence of intrinsic structure. Increasing substitution with native flours generally reduced viscosity, except for Weeping Grass, which was less affected and maintained stable viscosity. High-temperature rapid visco analysis revealed distinct thermal responses: most grains showed reduced final viscosity at elevated temperatures, whereas Weeping Grass maintained relatively high final viscosity across most temperatures. These findings demonstrate the diverse functional behaviour of Australian native grains and provide insights into their potential application in gluten-free and composite flour systems.

## Introduction

Among the physicochemical properties of flours from cereal grains and other sources, pasting behaviour is particularly important, as it influences the development of viscosity during heating and cooling in the presence of water. The Rapid Visco Analyser (RVA) is widely used to characterise these changes and provides viscosity profiles that are commonly interpreted as reflecting starch gelatinisation, granule stability, and molecular reassociation [1, 2]. Key RVA parameters, including peak viscosity (PV), trough viscosity (TV), breakdown viscosity (BDV), final viscosity (FV) and setback viscosity (SBV), describe distinct stages of pasting behaviour, including viscosity development, structural stability under thermal and shear conditions, and changes during cooling. Together, these parameters help evaluate flour functionality in applications such as baking, extrusion, and thickening of foods such as sauces, soups, gravies, and starch-based dispersions [3, 4].

In addition to the primary contribution of starch to the pasting properties of flours, other compositional and structural factors of flours can influence RVA profiles. Non-starch components, including proteins, lipids, and dietary fibre, can influence starch hydration and swelling by competing for water or restricting starch granule expansion [5, 6]. This effect is particularly evident in wholegrain flours, in which bran particles and cell wall materials can limit starch gelatinisation and are often associated with lower viscosity compared to refined flours. Therefore, interpretation of RVA results in wholegrain systems requires consideration of both compositional and structural factors [7, 8]. In addition to composition, the physical organisation of these components within the endosperm influences water accessibility and the behaviour of starch granules during heating, which contributes to differences in pasting behaviour [9, 10].

Flour particle size distribution is an important determinant of pasting behaviour, as it influences water accessibility, starch exposure, and interactions among flour components. Even under identical milling conditions and equipment settings, different grain species generally produce different proportions of coarse, medium, and fine particles [11, 12]. Thus, this variation in particle size distribution arises from intrinsic grain structure, which influences kernel fracture behaviour and particle formation during grinding [13].

In wheat, variation in flour particle size distribution between grains of different genotypes or plant growth environments is largely attributed to differences in endosperm hardness, which is related to the strength of intercomponent adhesion, particularly the interactions between starch granules and the surrounding protein matrix. It is also influenced by grain characteristics such as endosperm development, with poorly developed kernels generally producing fewer fine particles and more coarse fragments during milling [14, 15]. The grain moisture content strongly influences flour quality and milling yield. Grains with sufficient moisture tend to exhibit lower brittleness in the bran and endosperm, allowing more efficient separation during milling and a more consistent particle size distribution in the flour. In contrast, drier grains are more brittle, leading to greater fragmentation and a higher proportion of fine particles, including increased bran carryover into the flour, which can reduce flour quality and milling performance [15–17].

Water absorption capacity (WAC) is an important functional property that affects flour behaviour during hydration and thermal processing and is closely linked to pasting properties. It reflects the ability of flour components to absorb and retain water, thereby influencing starch gelatinisation, viscosity development, and final product texture [18, 19]. WAC depends not only on starch characteristics but also on protein, dietary fibre, and other non-starch components that compete for water or modify hydration behaviour, which affect water availability for starch swelling and gelatinisation. In this context, particle size plays a key role in determining water uptake and pasting behaviour. Finer particles generally exhibit faster water absorption due to increased surface area and improved water penetration; however, their effect on viscosity is not consistent, as starch damage and structural disruption may reduce viscosity development. In contrast, coarser particles may slow water uptake and limit viscosity development by restricting water penetration within the suspension. These effects are influenced by grain type and intrinsic structural characteristics [12, 20–22].

Among emerging agricultural resources, Australian native grain species have attracted increasing attention because of their adaptation to harsh environmental conditions, nutritional properties of the grain, and potential use in gluten-free and composite flour products. However, despite growing interest in these grains, their functional behaviour during processing remains poorly understood, particularly in relation to flour pasting behaviour [23, 24].

Flour functionality is also influenced by flour blending, which is commonly used to balance processing performance and nutritional value. Blending wheat flour with gluten-free flours can modify pasting behaviour due to differences in starch composition, protein functionality, and fibre content [25–27]. Higher substitution levels are often associated with changes in viscosity parameters, including reductions in PV and alterations in BDV and SBV [28, 29].

Thermal processing strongly affects starch functionality, as higher temperatures promote gelatinisation, breakdown of granules, and molecular changes, which in turn influence viscosity and final product structure. Elevated temperatures used in industrial processes such as extrusion, high-temperature cooking, and sterilisation can cause starch to behave differently from conventional pasting conditions [30–33].

This study aimed to investigate the effects of particle size, flour blending, and elevated processing temperatures on the pasting behaviour of four Australian native grains: Button Grass, Curly Mitchell Grass, Native Millet, and Weeping Grass. Particle size distribution was determined by quantifying the proportions of fine, medium, and coarse fractions obtained through sieving. RVA pasting properties were then analysed for each fraction and compared to evaluate the effect of particle size. Composite flours were prepared by blending wholegrain native flours with refined wheat flour at substitution levels of 0, 5, 10, 20, and 30%, and their pasting properties were assessed. In addition, high-temperature RVA analysis was conducted on the wholegrain flours of each native grain to emulate processing conditions that involve higher levels of heat.

## Materials and Methods

### Grain Materials

Mature seeds of four Australian native grain species—Button Grass (*Dactyloctenium radulans*), Curly Mitchell Grass (*Astrebla lappacea*), Native Millet (*Panicum decompositum*), and Weeping Grass (*Microlaena stipoides*) —were the main grains investigated. Wheat (*Triticum aestivum* cv. Kord) and brown teff (*Eragrostis tef*, a gluten-free grain) were included as reference grains. In addition, refined wheat flour (*Triticum aestivum* cv. Lancer) was used for composite flour preparation.

Button Grass and Curly Mitchell Grass seeds were obtained from the same location in Thallon, Queensland, Australia. Native Millet seeds were sourced from the University of Sydney Plant Breeding Institute and L’lara Farm in Narrabri, New South Wales, Australia. Weeping Grass seeds were supplied by a commercial native food producer (Creative Native Food Service Co, Australia), harvested near Armidale, New South Wales. Brown teff grain was purchased commercially from an Australian supplier (Aussie Coffee Roasters, Australia) and, according to the supplier, was grown in New South Wales. Refined wheat flour was purchased from Provenance Flour and Malt (Australia).

### Flour Preparation and Particle Size Fractionation

Whole grains were milled under identical conditions with a Laboratory Mill 3100 (Perten, Australia) fitted with a 500 µm screen to obtain wholemeal flour. A 100-g flour sample was fractionated by dry sieving using a test sieve shaker (Endecotts EFL 2 Mk3, UK). A sieve stack with aperture sizes of 355 µm and 180 µm was used to separate the flour into >355 µm, 180– 355 µm, and <180 µm fractions. Samples were sieved for 10 min under constant shaking conditions [12]. Following sieving, the material retained on each sieve was collected and weighed. Particle size distribution was expressed as the proportion (%) of the total sample mass in each fraction [34].

### Composite Flour Preparation

Composite flours were prepared by blending native grain flours with refined wheat flour (cv. Lancer) at substitution levels of 0, 5, 10, 20, and 30% (w/w), with the 0% sample consisting of wheat flour only and serving as the control. For each blend, the required amounts of native grain and wheat flours were weighed to obtain the desired ratios. Samples were transferred to 50-mL tubes and mixed using an Intelli-Mixer RM-2 (John Morris, Australia) using mode F6 (30 rpm) for 20 min to ensure uniform distribution of components. All blends were prepared and analysed on the same day.

### Moisture Content Measurement

Moisture content was determined using the AACC 44–15.02 method [35]. Approximately 2 g of sample was weighed into pre-weighed aluminium dishes and dried at 135 ± 1°C for 2 h. Samples were cooled in a desiccator to room temperature before reweighing. Moisture content was calculated from the weight loss during drying.

### Rapid Visco Analysis (RVA)

Pasting properties of flour samples were evaluated using a Rapid Visco Analyser (RVA 4800, Perten Instruments, Australia) operated with ThermoCline for Windows (TCW) version 3.17 software. For each measurement, 3.5 g of flour, adjusted to a 14% moisture basis based on measured moisture content, was weighed into RVA canisters, followed by the addition of 25 g of Milli-Q water.

All samples were analysed using a standard pasting profile based on AACC International Method 76-21.02 [36]. The specific time–temperature–speed conditions for each pasting program are summarised in Table 1. The standard profile included equilibration at 50 °C, heating to 95 °C, a holding stage, cooling to 50 °C, and a final holding stage. High-temperature profiles were modified to reach maximum temperatures of 120, 130, and 140 °C before cooling, as detailed in Table 1.

**Table 1.**
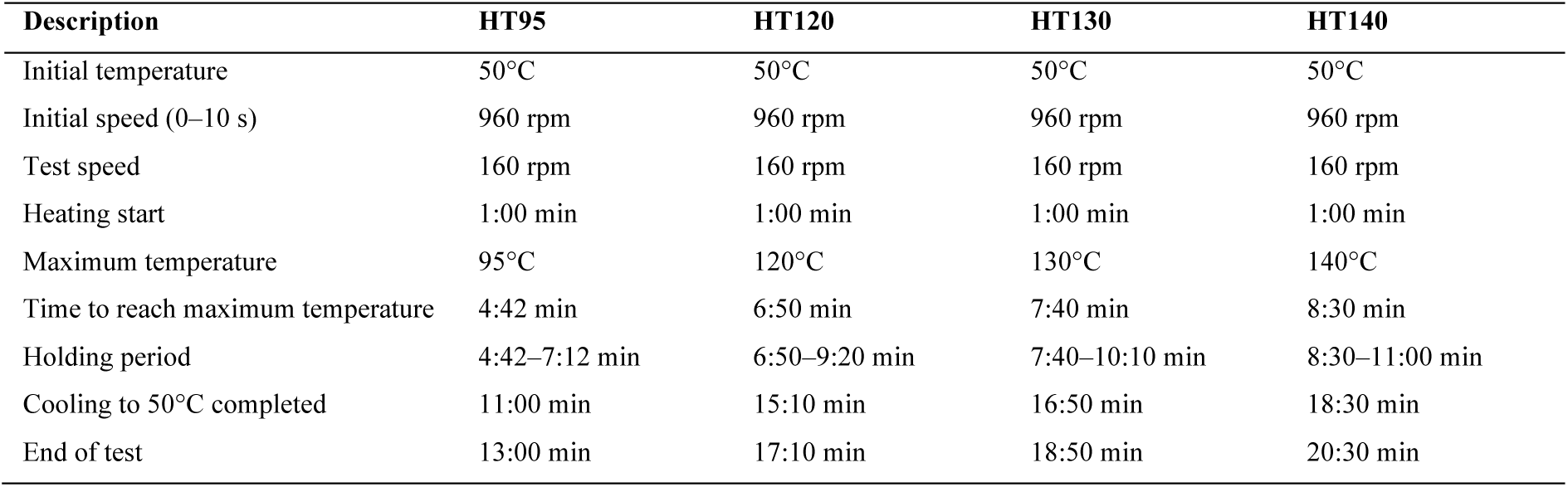
Time–temperature–speed profiles for the standard AACC method (95 °C) and modified high-temperature (120, 130, and 140 °C) RVA pasting programs. HT95, HT120, HT130, and HT140 denote RVA standard and high-temperature profiles with maximum temperatures of 95, 120, 130, and 140°C, respectively.

### Water Absorption Capacity

WAC was determined as described by Abd Elmoneim et al. (2010). Flour (1.0 g) was added to a pre-weighed 15-mL conical centrifuge tube, and the combined weight was recorded as W₁. Distilled water (10 mL) was added, and the mixture vortexed for 1 min, then left to stand at room temperature (25 ± 2 °C) for 30 min. The samples were then centrifuged at 4,000 × *g* for 25 min, after which the supernatant was decanted. The tube containing the hydrated flour was reweighed (W₂), and WAC was calculated using the following equation:

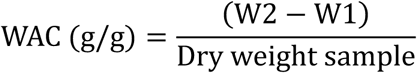

### Statistical Analysis

All RVA measurements were performed in duplicate using independently prepared test portions from the same flour sample, and results are expressed as mean ± standard deviation [38]. One-way analysis of variance (ANOVA) was used to assess differences among particle size fractions within each grain and among substitution levels for RVA parameters, as well as among grains for WAC, where applicable. Where significant differences were detected for WAC, Tukey’s *post hoc* multiple comparisons test was applied. Statistical significance was set at *p* < 0.05. All analyses were performed using GraphPad Prism (version 10.2.1, GraphPad Software, USA).

## Results and Discussion

### Particle Size Distribution of Native and Reference Grain Flours

The particle size distribution of the whole-grain milled flours showed clear differences among the native grain species and the reference grains (Figure 1). Button Grass and Weeping Grass flours had a high proportion of fine particles (<180 µm), representing 58.5% and 70.6% of the total, respectively. In contrast, Native Millet flour was mainly composed of medium-sized particles (70.5%), whereas Curly Mitchell Grass flour showed a more balanced distribution between medium (55.1%) and fine (41.6%) fractions.

**Figure 1.**
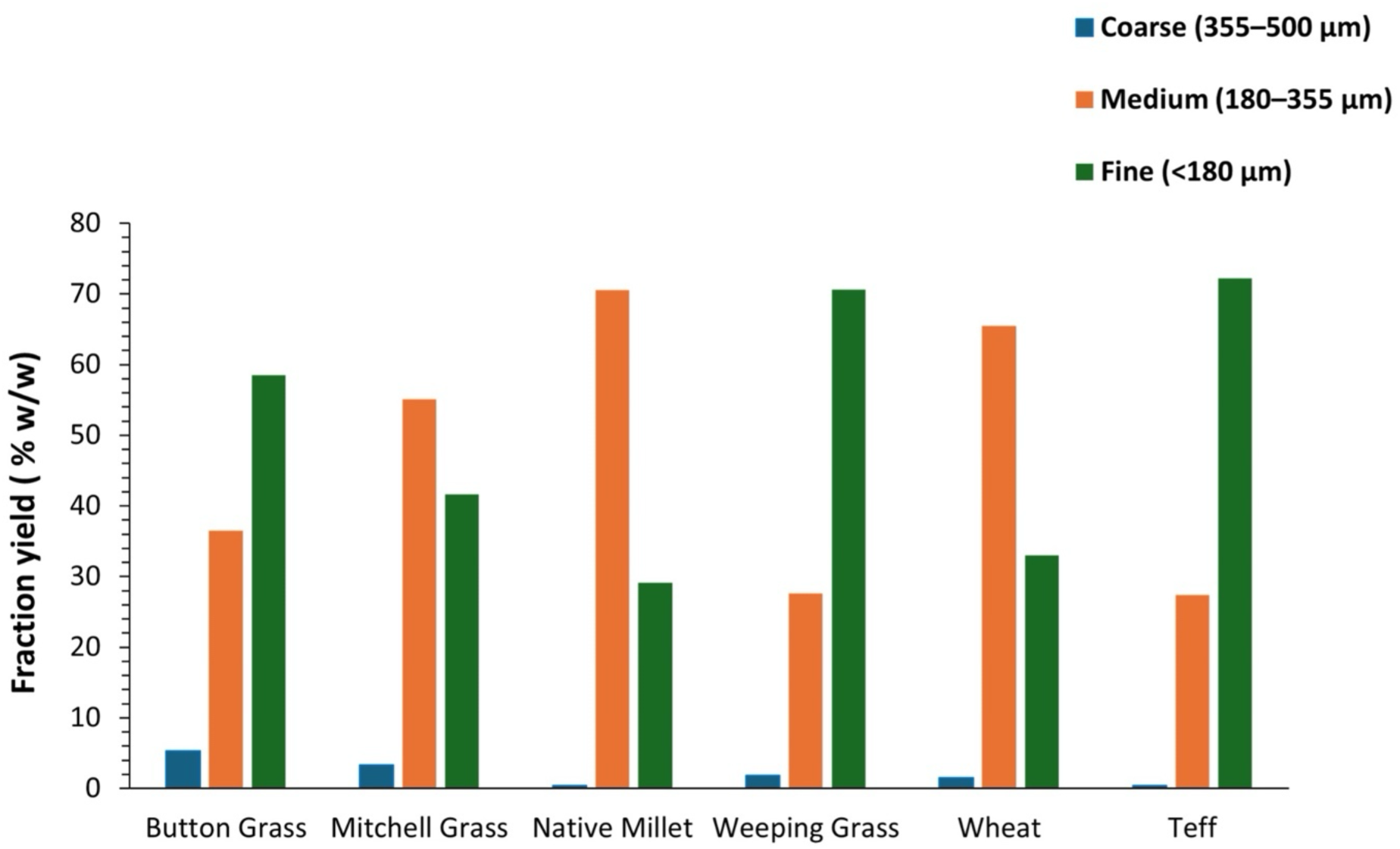
Particle size distribution of milled flours from Australian native grains and reference grains, expressed as fraction yield (% w/w) across coarse (355–500 µm), medium (180–355 µm), and fine (<180 µm) fractions. Whole grains were milled into wholemeal flour using a Laboratory Mill 3100 (Perten, Australia) fitted with a 0.5 mm screen before sieving.

Among the reference grains, wheat flour was primarily composed of medium-sized particles (65.5%) with a lower proportion of fines, while teff showed a predominance of fine particles (72.2%) with a negligible coarse fraction (0.4%).

Differences in flour particle size distribution under comparable milling conditions are common in cereal systems and are influenced by kernel properties such as hardness, endosperm structure, and texture, as well as fracture behaviour during milling [14, 39–41]. In cereal endosperm, starch granules are embedded within a protein matrix, and this structural organisation contributes to the physical properties of the grain [42, 43]. Grains with a more compact endosperm generally produce larger particles, whereas less compact structures tend to produce finer fractions during milling [41]. These differences reflect variation in the mechanical strength of the endosperm matrix and the adhesion between starch granules and surrounding components, as widely reported in cereals, particularly wheat [15, 44, 45]. Native Millet showed a higher proportion of medium-sized particles, which may relate to a more densely packed starch structure, as suggested by previous microscopy observations using Lugol’s iodine staining [46]. In contrast, Button Grass and Curly Mitchell Grass showed less densely packed starch granule arrangements (ibid.) and exhibited higher proportions of fine particles (Figure 1), which may indicate greater susceptibility to fragmentation. Such relationships between starch structure, kernel properties, and milling behaviour are established in cereal grains and are consistent with the patterns observed here [14, 47].

Compared with wheat, the flour of which was dominated by the medium fraction, the native grains displayed a broader distribution of particles across fine, medium, and coarse fractions. In modern processing of commercial cereals, milling equipment and conditions allow a high degree of control and consistency in the particle size distribution of flour fractions [48]. In contrast, wild cereal species often show greater structural variability, which can lead to less uniform milling behaviour [49]. This variation highlights the importance of grain-specific structural and compositional properties in the design of processing strategies.

### Effect of Particle Size on Pasting Behaviour of Native and Reference Grain Flours

The pasting profiles of the native and reference grains demonstrated clear but grain-specific responses to particle size classes, as observed in the RVA curves (Figure 2) and supported by the standardised heatmap (Figure 3). All samples exhibited typical starch pasting behaviour, including viscosity increase during gelatinisation, followed by breakdown and subsequent viscosity development during cooling. However, these changes were noticeably different depending on the grain type and particle size.

**Figure 2.**
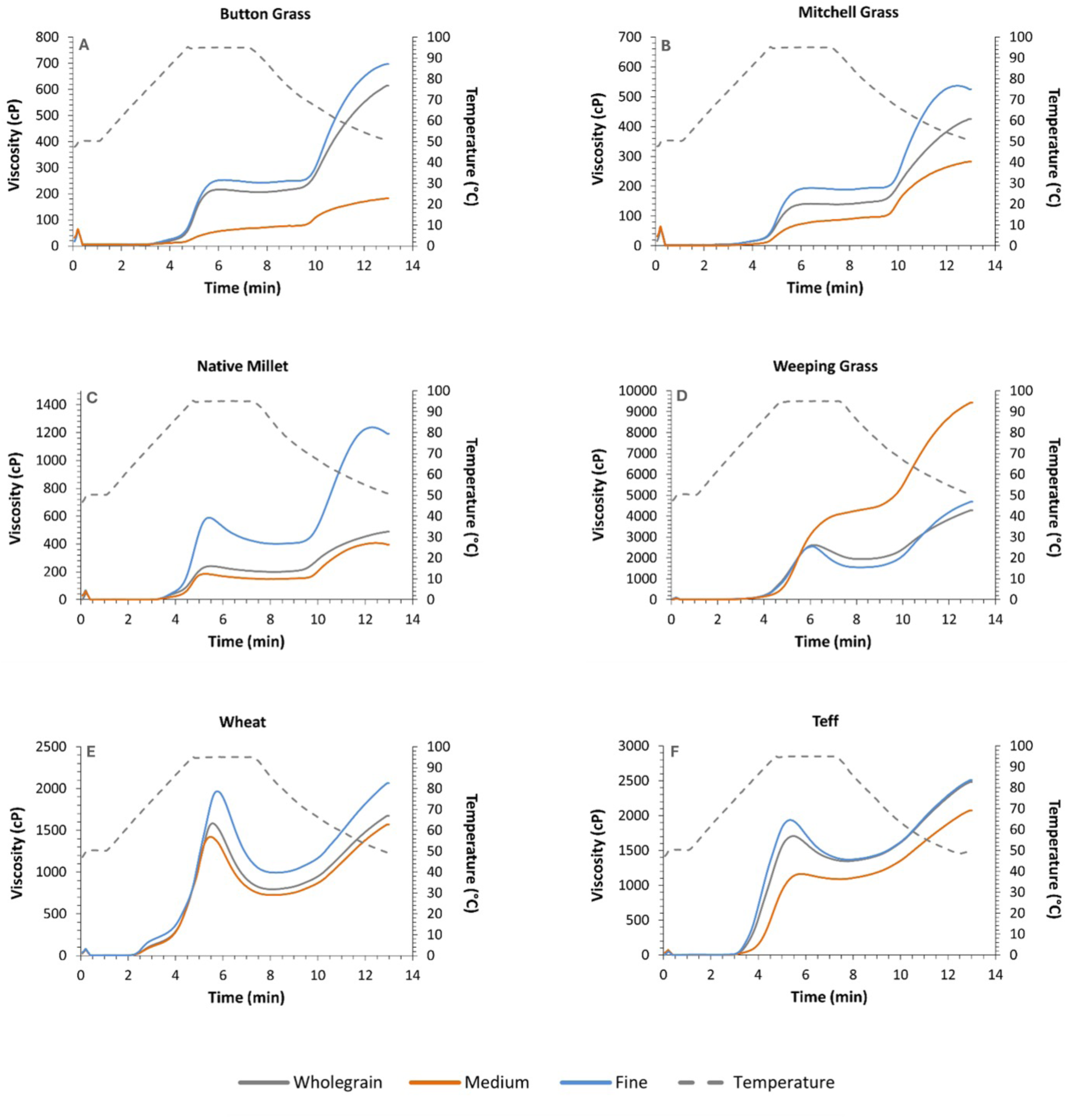
Pasting profiles of native and reference grain flours with different particle size ranges (wholegrain, medium: 180–355 µm, and fine: <180 µm), measured using an RVA 4800. Viscosity (cP) changes over time (min) were determined, with the temperature profile (°C) shown as a dashed line. Panels (A–F) correspond to Button Grass, Curly Mitchell Grass, Native Millet, Weeping Grass, wheat, and teff, respectively. Note that the viscosity scale on the y-axis varies between panels to show differences among samples more clearly, as some flours (e.g., Weeping Grass) reached much higher viscosity values than others. Variations in pasting parameters (e.g., PV and FV) reflect not only particle-size differences but also compositional factors, such as the amylose: amylopectin ratio. Detailed experimental conditions are provided in the Materials and Methods section.

**Figure 3.**
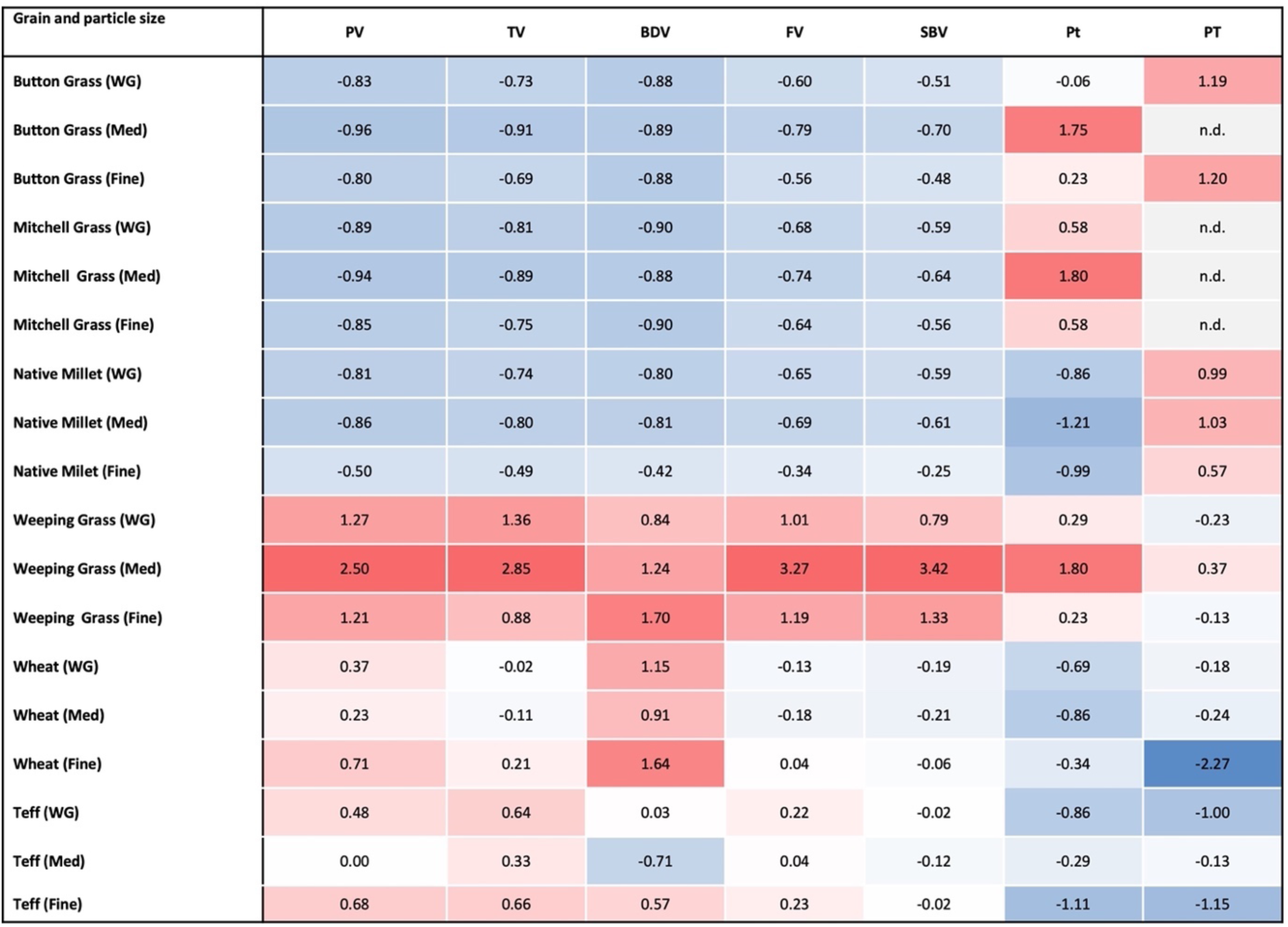
Heatmap of standardised pasting properties (peak viscosity – PV, trough viscosity – TV, breakdown viscosity – BDV, final viscosity – FV, setback viscosity – SBV, peak time – Pt, and pasting temperature – PT) of native grains, wheat, and teff across different particle sizes. WG, Med, and Fine denote wholegrain, medium, and fine particle sizes, respectively. Values are expressed as z-scores to allow comparison among variables with different scales, where 0 represents the overall mean for each parameter and positive or negative values indicate the number of standard deviations above or below the mean, respectively. For example, a value of 2.85 indicates that the measured parameter was 2.85 standard deviations higher than the overall mean for that parameter. Red and blue indicate relatively higher and lower values, respectively. Cells labelled “n.d.” indicate samples for which data could not be obtained because of instrument-related limitations during analysis.

### Peak Viscosity

Among all samples, Weeping Grass at medium particle size exhibited the highest PV and the highest standardised value in the heatmap analysis (Figures 2D and 3), with a higher PV than its wholegrain and fine fractions (*p* < 0.05). This behaviour may be partly related to the relatively high WAC of Weeping Grass observed (Figure 4), which was among the highest recorded values. Higher water absorption may increase water availability during heating and promote granule swelling, which may result in higher paste viscosity [50]. This effect may be further supported by the high β-glucan content of Weeping Grass [51]. Cereal β-glucans possess high water-binding capacity and are known to increase viscosity in cereal systems [52, 53].

**Figure 4.**
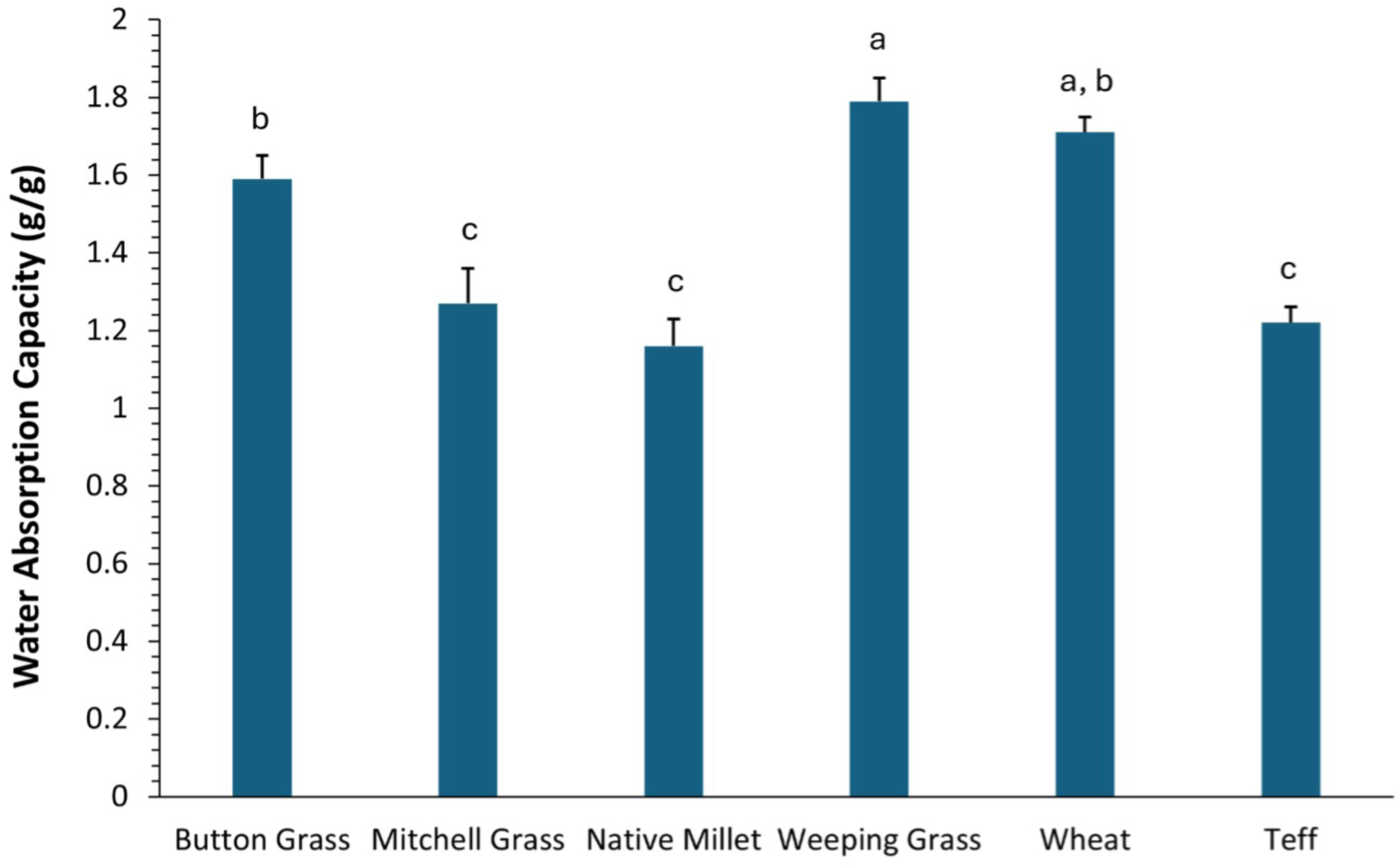
Water absorption capacity (g water/g sample) of native grains (Button Grass, Curly Mitchell Grass, Native Millet, and Weeping Grass) and reference grains (wheat and teff). All samples were milled into wholemeal flour from the same batch. Values represent the means of duplicate measurements, with error bars indicating standard deviation [38]. Means with different letters are significantly different (*p* < 0.05), as determined by Tukey’s multiple comparisons test following one-way ANOVA.

In contrast to Weeping Grass, Native Millet displayed its highest PV in the fine fraction, which was a higher value than for both the wholegrain and medium fractions (*p* < 0.05). This suggests that finer milling increased starch exposure in Native Millet, which enhanced water penetration and promoted greater granule swelling during heating [54, 55]. The lower PV observed in the wholegrain fraction may reflect the presence of bran and fibre components that limited starch swelling and reduced viscosity. This interpretation is consistent with previous findings for Native Millet, where wholemeal flour exhibited lower pasting ability due to higher fibre content and a greater bran-to-endosperm proportion [23]. This behaviour may also reflect the relatively low WAC of Native Millet wholegrain flour observed here (Figure 4), which may limit starch–water interactions compared with grains that exhibited higher WAC [18, 19].

Similar to Native Millet, wheat showed higher PV in the fine fraction than both the wholegrain and medium fractions (*p* < 0.05), while the difference between the wholegrain and medium fractions was not significant. In contrast, teff showed a stronger response to particle size, with the fine fraction higher in PV than both the wholegrain and medium fractions (*p* < 0.05). The wholegrain fraction also had higher PV than the medium fraction (*p* < 0.01). This behaviour is particularly relevant for gluten-free product development, where starch swelling and viscosity development are critical for providing structure in the absence of gluten [56, 57].

Button Grass and Curly Mitchell Grass showed limited sensitivity to particle size, and standardised PV values remained low across all fractions (Figures 2A, 2B and 3). This behaviour may be influenced by compositional and structural factors, including the relatively high protein content of Curly Mitchell Grass [51], as protein can restrict starch swelling by limiting water availability and may reduce the effect of particle size on pasting behaviour [58]. These results indicate that particle size had a relatively small effect on swelling behaviour, while intrinsic structural characteristics of these grains played a more dominant role. Previous studies on wild cereal species have reported differences in starch granule morphology, endosperm organisation, and greater retention of cell wall material at maturity, which may further restrict starch swelling and contribute to a limited pasting response [59].

For Button Grass, the limited sensitivity to particle size may relate to its relatively high WAC (Figure 4). A portion of the absorbed water may associate with fibre and protein components, which reduces water availability for starch gelatinisation. Previous studies show that non-starch polysaccharides can limit water access to starch granules and restrict swelling during pasting, which may contribute to lower viscosity despite relatively high water uptake [60–62].

### Breakdown Viscosity and Paste Stability

Breakdown viscosity, which indicates the degree of structural disruption of swollen starch granules under thermal and shear conditions [63, 64], generally followed trends similar to PV in some samples, particularly the fine fractions of Native Millet, wheat, and teff, where higher PV was associated with greater BDV. This trend was most pronounced in Native Millet, where the fine fraction had higher BDV than both the wholegrain and medium fractions (*p* < 0.0001). Teff showed differences across particle sizes, with BDV lowest in the medium fraction, higher in the wholegrain, and highest in the fine fraction (*p* = 0.0005). In wheat, the fine fraction had higher BDV than both the wholegrain and medium fractions (*p* = 0.0068). These results indicate that greater starch granule swelling was associated with increased susceptibility to structural disruption during heating [65]. However, this relationship was not consistent across all grains or particle size fractions, as BDV depends on both PV and TV, and variation in TV influenced the extent of breakdown.

The relationship between PV and BDV was not consistent across all samples. In Button Grass and Curly Mitchell Grass, the small decline from PV to TV indicated greater paste stability during the holding phase, which resulted in relatively low BDV values. The relatively small decline in viscosity between PV and TV, compared with the more pronounced drop observed in wheat and teff, suggests limited granule collapse during holding. This behaviour may relate to restricted granule swelling and lower susceptibility to shear-induced disintegration [64–66].

Weeping Grass showed a distinct pasting profile, particularly in the medium fraction, which had high PV and a moderate decline from PV to TV, indicating an intermediate level of breakdown compared with other grains. This suggests that, despite some structural disruption during heating, the paste retained a degree of stability under thermal and shear conditions.

### Final Viscosity and Retrogradation Behaviour

Final viscosity represents the value measured at the end of the RVA test after cooling and reflects molecular reassociation under the defined test conditions. As viscosity development depends on time, cooling, and holding conditions, further increases may occur if the test duration is extended [67, 68]. The FV data showed clear grain-specific differences in response to particle size (Figure 2). In wheat, the fine fraction exhibited the highest FV, followed by the wholegrain and medium fractions. In teff, both the fine and wholegrain fractions showed similarly high FV values, whereas the medium fraction was lower, indicating a non-linear response to particle size.

The behaviour of Weeping Grass in relation to FV was particularly notable. The medium fraction showed higher FV than both the wholegrain and fine fractions (*p* < 0.001), while no significant difference was observed between the wholegrain and fine samples (Figure 2D). This indicates that the medium particle size promoted greater molecular reassociation during cooling and stronger gel formation [69, 70]. A similar trend was observed for SBV, where the medium fraction also showed higher values than both the wholegrain and fine fractions (*p* < 0.01).

Strong viscosity development during cooling may be advantageous in gluten-free applications, where starch gel formation plays an important role in improving structure formation and crumb stability in the absence of gluten [56, 71]. This behaviour may be linked to the high β-glucan content of Weeping Grass [51], which contributes to increased viscosity during both heating and cooling, including enhanced gel formation during cooling [72, 73]. However, strong viscosity development during cooling may also have implications for staling, although this is more directly reflected by setback viscosity rather than final viscosity [74, 75].

In Curly Mitchell Grass, FV varied with particle size, with the highest value in the fine fraction, followed by the wholegrain and medium fractions. SBV showed a similar pattern, which indicates greater molecular reassociation and gel formation in the fine fraction.

In Native Millet, FV was affected by particle size with the value for the fine fraction exceeding those for both the wholegrain and medium fractions (*p* < 0.01), while no difference was found between the latter two. The higher FV in the fine fraction probably reflects increased starch accessibility, which promotes molecular reassociation during cooling. In contrast, the lower FV in the wholegrain fraction may result from the presence of bran and fibre components that limit starch swelling and reduce water availability [8, 76, 77]. Although amylose content can influence viscosity development during cooling, previous work has shown that the wholegrain of this millet has a relatively high amylose content; however, as amylose content was not determined for the individual fractions in this study, it cannot be used to explain the observed differences in FV between fractions.

Button Grass showed relatively low FV values, with the medium fraction lower than both the wholegrain and fine fractions (*p* ≤ 0.0001). This pattern indicates limited viscosity development during cooling, which suggests reduced starch reassociation compared with the other grains.

### Effect of Native Grain Substitution on Pasting Behaviour of Wheat Composite Flours

Overall, viscosity tended to decrease with increasing substitution of refined wheat flour with native grain flour (Figure 5). However, this pattern varied between species, with a clearer reduction observed in Button Grass, Mitchell Grass, and Native Millet, whereas Weeping Grass showed little change across substitution levels.

**Figure 5.**
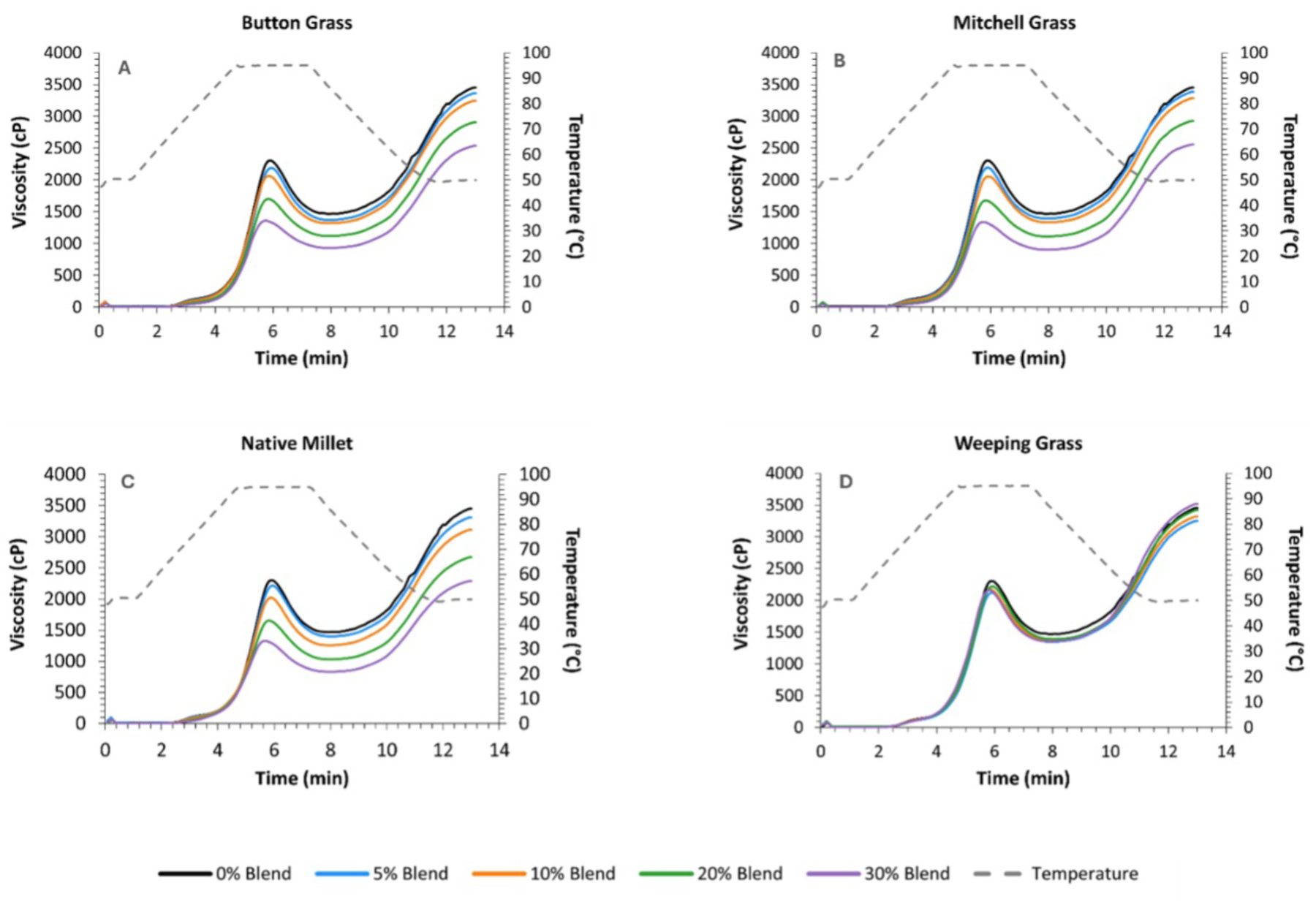
RVA pasting profiles of Lancer wheat endosperm flour (0% blend) and composite flours containing 5–30% native grains blended with Lancer wheat. Viscosity (cP) is shown over time (min), with the temperature profile (°C) indicated by the dashed line. Panels (A–D) represent Button Grass, Curly Mitchell Grass, Native Millet, and Weeping Grass, respectively. Experimental details are provided in the Materials and Methods section.

The trend above was further reflected in Button Grass, Curly Mitchell Grass, and Native Millet, where PV, TV, and FV decreased progressively with increasing substitution levels. Across all three grains, PV and FV decreased at higher substitution levels (20 and 30%) compared with the control (*p* < 0.05) and tended to decline further at 30%. Trough viscosity showed a similar downward pattern across the samples, and SBV generally decreased as substitution increased, which suggests a lower tendency for starch reassociation during cooling. This behaviour is likely due to changes in starch functionality rather than a simple reduction in starch content. Although both wheat and native grains contain starch, differences in composition and structure, along with the presence of fibre and protein, can limit starch hydration and swelling, reducing its contribution to viscosity during heating [76, 78, 79]. From an application perspective, the consistent reduction in PV and FV with flour substitution suggests a potential limitation in the use of these blends for food products that require strong viscosity development and structure formation [2, 30].

In contrast to the behaviour above, increasing Weeping Grass substitution had little effect on PV, FV, and SBV, which suggests more stable starch behaviour within the composite system. This may be partly related to the relatively high WAC of Weeping Grass observed in the present study, which could support starch hydration during heating. In addition, its β-glucan content may also play a role in maintaining viscosity, as β-glucans are known to enhance water binding and increase paste viscosity in cereal systems [53, 80, 81].

Overall, the effects of blending were not uniform across the native grains. Button Grass, Curly Mitchell Grass, and Native Millet showed consistent reductions in viscosity with increasing substitution, whereas Weeping Grass was less affected and showed more stable pasting behaviour during both heating and cooling. These differences highlight the importance of grain-specific compositional and functional characteristics in determining the pasting behaviour of wheat-based composite flours.

### Effect of Maximum Temperature on RVA Pasting Behaviour of Flour Samples

As the maximum temperature increased from 95 to 140 °C, the RVA pasting profiles changed noticeably, with clear differences between the grains (Figure 6). At 95 °C, all samples showed the typical pattern, where viscosity increased during heating, dropped during the holding stage, and rose again during cooling. As the temperature increased, both the level and shape of the viscosity curves changed, although the extent and direction of these changes differed between the species.

**Figure 6.**
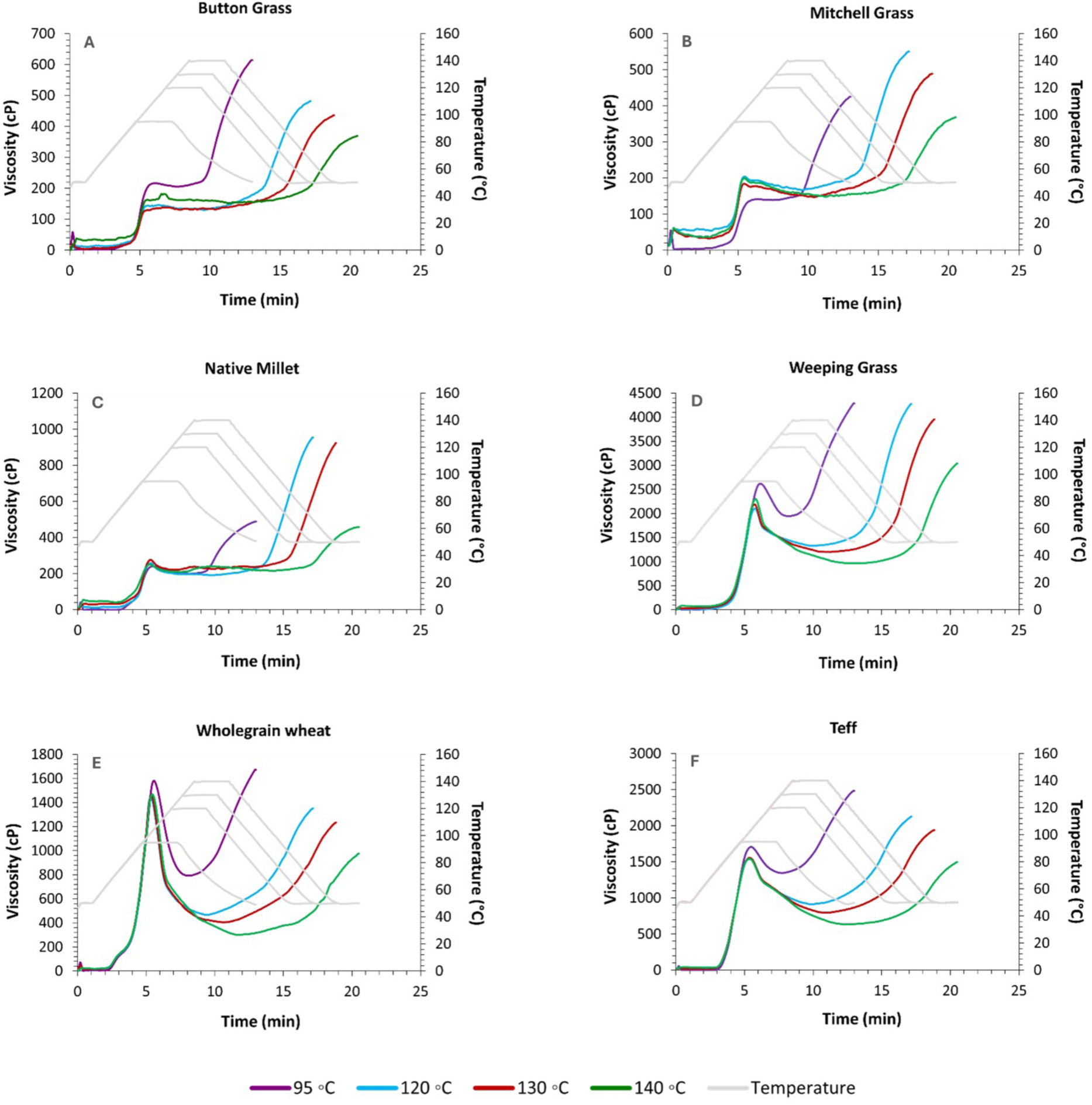
RVA pasting profiles of flour samples at different maximum temperatures (95, 120, 130, and 140 °C). Viscosity (cP) is presented as a function of time (min), with the corresponding temperature profile (°C) shown in grey. Panels (A–D) correspond to Button Grass, Curly Mitchell Grass, Native Millet, and Weeping Grass, respectively. The curves illustrate the influence of increasing temperature on pasting characteristics, including PV, BDV, and FV. Experimental details are provided in the Materials and Methods section.

For Button Grass, PV decreased with increasing temperature up to 130 °C, followed by a slight increase at 140 °C (Figure 6A). Final viscosity also tended to decrease with increasing temperature, suggesting reduced viscosity development during cooling. This response may reflect higher thermal disruption of starch granules at elevated temperatures, which can weaken paste structure and reduce viscosity. The increase in PV at 140 °C may indicate a shift in the balance between granule swelling and structural breakdown under more severe heating conditions [82, 83].

Curly Mitchell Grass showed a different pattern with temperature (Figure 6B), where PV increased up to 120 °C and then remained relatively stable at higher temperatures. Final viscosity increased from 95 to 120 °C, where it reached its highest value, and then decreased at 130 and 140 °C, indicating reduced viscosity development during cooling at higher temperatures. This behaviour may be related to improved water access to starch granules and greater release of starch components at 120 °C, which supported viscosity development. At higher temperatures, swollen granules may have lost structural integrity and become more prone to breakdown, which weakened the paste structure and reduced viscosity during cooling [19, 65, 83].

In Native Millet, PV showed little change across the temperature range (Figure 6C). In contrast, final viscosity increased from 95 to 120 and 130 °C, before dropping sharply at 140 °C, indicating reduced viscosity development during cooling at higher temperatures. This sharper decline implies greater sensitivity to high temperature. This pattern suggests that moderate temperatures support starch swelling and viscosity development, whereas at 140 °C, greater structural disruption of swollen granules likely limits viscosity during cooling [19, 82].

In contrast to Native Millet, Weeping Grass showed a markedly different response to temperature (Figure 6D). Peak viscosity was highest at 95 °C and decreased at higher temperatures. However, FV remained relatively high across all conditions, increasing up to 130 °C before declining at 140 °C, which indicates strong viscosity development during cooling. This behaviour shows that, despite the lower PV at higher temperatures, the paste still developed a strong structure during cooling. This suggests that, despite lower PV at higher temperatures, Weeping Grass retained a strong capacity for structure formation during cooling. The higher FV may also indicate greater gel firmness and a stronger tendency for retrogradation, which could be relevant to texture development in starch-based products [19, 84].

Wheat and teff showed similar behaviour, with PV decreasing from 95 to 120 °C and then remaining relatively stable at higher temperatures. In contrast, FV decreased progressively from 95 to 140 °C in both grains, indicating reduced viscosity development during cooling at higher temperatures. This behaviour may be related to increased thermal disruption of starch granules at elevated temperatures, which limits further viscosity development during heating and reduces the ability of starch components to reassociate during cooling [82, 85].

## Conclusion

This study provides insights into the structure–function relationships of Australian native grain flours by examining the effects of particle size, flour blending, and high temperature on pasting behaviour. The results reveal grain-specific responses that reflect differences in composition, structure, and water interactions.

Particle size affected pasting behaviour differently among the grains. In Native Millet, finer particles were associated with higher overall viscosity, likely due to increased starch exposure and improved water interaction. In contrast, Button Grass showed a pronounced reduction in viscosity in the medium fraction compared to both wholegrain and fine fractions, indicating a strong but non-linear sensitivity to particle size. Curly Mitchell Grass showed a more moderate response, suggesting that intrinsic structural and compositional features, such as starch granule accessibility, fibre content, and starch–protein interactions, influenced pasting behaviour by affecting water absorption, granule swelling, and viscosity development during heating [86].

Weeping Grass flour had its highest viscosity at medium particle size rather than in the fine fraction, particularly in terms of final viscosity, highlighting that the relationship between particle size and viscosity is grain-specific and not always linear.

Flour blending reduced viscosity in most composite systems, likely due to starch dilution and competition for water from non-starch components such as protein and fibre. This effect was comparatively limited for blends up to 10% of native flour addition. In contrast, Weeping Grass maintained relatively stable viscosity under all substitution levels, suggesting greater stability during heating and cooling. This behaviour may be associated with its water absorption capacity and β-glucan content, which contribute to viscosity development.

The high-temperature RVA results further highlight differences in thermal response among the grains. Button Grass showed an overall reduction in final viscosity as temperature increased, similar to wholegrain wheat and teff, which reflects lower viscosity development during cooling at higher temperatures. Curly Mitchell Grass showed greater viscosity development at 120 °C, with lower values at higher temperatures. In contrast, Weeping Grass maintained relatively high FV across the temperature range, which suggests a stronger capacity for structure formation during cooling. Native Millet also exhibited elevated FV at certain temperatures; however, its response was more variable and non-linear, indicating a different underlying structural behaviour. Final viscosity provided insight into the functional behaviour of the grains and their potential performance in food systems. Higher FV, particularly in Weeping Grass, suggests greater viscosity development and gel formation during cooling, which may be advantageous in applications that require structural integrity, such as porridges, gels, or thickened food products.

Stronger molecular reassociation may also increase the tendency for gel firming during storage. In contrast, Button Grass showed lower FV, suggesting limited gel formation and reduced structure development during cooling. This behaviour may be more suitable for applications that require a softer or less structured gel texture, such as sauces and soups. Native Millet and Curly Mitchell Grass showed intermediate behaviour, which suggests potential use in formulations that require a balance between high and low-viscosity systems.

Australian native grains show distinct functional properties and should not be considered direct equivalents to conventional cereals such as wheat and teff. As naturally gluten-free ingredients, each grain behaves differently and can be used to adjust texture, stability, and shelf life across a range of foods. These findings support the potential of native grains as functional ingredients in modern and gluten-free products and highlight the need for grain-specific processing strategies to optimise their performance. In addition, the growing interest in native grains may support the development of Indigenous-led enterprises and contribute to broader recognition of traditional knowledge and sustainable grain use.

## Acknowledgments

We gratefully acknowledge the Gamilaraay people for sharing their knowledge of native grains and traditional food systems. We particularly thank Kerrie Saunders for her generous guidance, support, and assistance with native grain collection and processing.

## Author contributions

**Farkhondeh Abedi:** Conceptualization, Methodology, Investigation, Formal analysis, Visualization, Writing – original draft.

**Ali Khoddami:** Supervision, Writing – review & editing.

**Claudia Keitel:** Supervision, Writing – review & editing.

**Thomas H. Roberts:** Conceptualization, Methodology, Project administration, Resources, Supervision, Writing – review & editing.

